# A polymer-physics view of peripheral chromatin: de Gennes’ self-similar carpet

**DOI:** 10.1101/2024.01.05.574343

**Authors:** Ozan S. Sarıyer, Aykut Erbaş

## Abstract

Using scaling arguments to model peripheral chromatin localized near the inner surface of the nuclear envelope (NE) as a flexible polymer chain, we discuss the structural properties of the peripheral chromatin composed of alternating lamin-associated domains (LADs) and inter-LADs. Modeling the attraction of LADs to NE by de Gennes’ self-similar carpet, which treats the chromatin layer as a polymer fractal, explains two major experimental observations: (*i*) The high density of chromatin close to the nuclear periphery decays to a constant density as the distance to the periphery increases. (*ii*) Due to the decreasing mesh size towards the nuclear periphery, the chromatin carpet inside NE excludes molecules (*via* non-specific interactions) above a threshold size that depends on the distance from the nuclear periphery.

## I. INTRODUCTION

Inside the highly crowded volume of the eukaryotic cell nucleus, chromatin is not randomly distributed [1– 7]. While euchromatin, which allows access to lineage-specific genes, is often observed away from the nuclear surfaces, heterochromatin, which is commonly associated with gene silencing, is localized near the nuclear surfaces [4–10]. This 3d organization scheme is also one of the epigenetic factors regulating the biological characteristics and health of a cell [11, 12]. Consistently, many genetic disorders with a terminal nature, such as various types of laminopathies [13–18] and some cancers [19, 20], manifest themselves as alterations in peripheral chromatin distribution. Understanding the origins of such distorted distribution patterns requires characterizing structural and conformational properties of peripheral chromatin localized near the inner surface of the nuclear envelope (NE).

Recent progress in sequence-based chromatin conformation capture technologies (e.g., Hi-C [21], ChIP [22], and DamID [23]) in addition to developments in high-resolution-microscopy (e.g., FISH [24]), have provided the most detailed structural properties of peripheral chromatin so far [7, 25]. These experiments report that peripheral chromosomes have many distinct lamina-associated domains (LADs), which interact with nuclear boundary components and have the characteristics of gene-inactive heterochromatin. These domains have sharply defined boundaries, and their lengths range between 0.1 Mb (mega-basepairs) to 10 Mb with a median length of about ∼ 0.5 Mb [7, 26–31]. LADs are connected by relatively longer strands of inter-LADs, which form chromatin loops protruding towards the nuclear interior (Fig. 1). Interestingly, unlike LADs, inter-LADs exhibit the characteristics of gene-active euchromatin [31– 34]. Consistently, experiments reported that large multiprotein complexes maintaining gene transcription (*e.g*., RNA polymerase II) and enzymes involved in DNA repair machinery (*e.g*., DNA glycosylase) interact less with LADs than with inter-LADs [7, 9, 26]. Relatedly, experiments probing chromosome structures with nanometer-sized dextran particles showed that large probe particles could not penetrate near the nuclear boundary [35], which suggests a denser (chromatin) environment with smaller mesh sizes at the nuclear periphery, in comparison to central nuclear space.

**FIG. 1.**
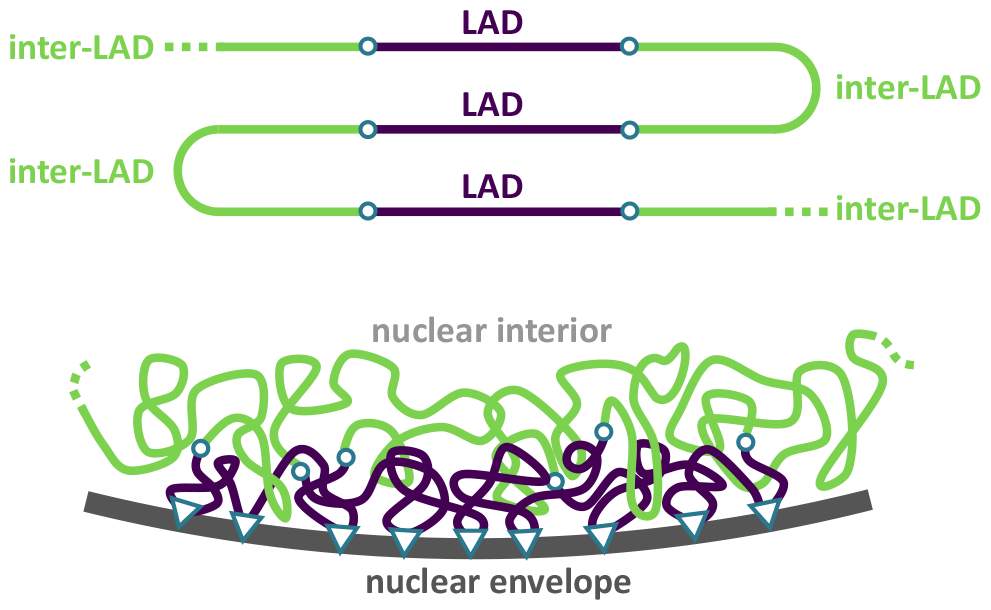
Authors’ view of peripheral chromatin. A polymer train of alternating LADs (purple) and inter-LADs (green) along a peripheral chromosome. Open circles indicate the sharp LAD/inter-LAD borders. Various proteins anchoring LADs to the nuclear boundary are represented with triangles. The real biological environment at the eukaryotic nuclear periphery may have a higher chromatin density than shown here.

LADs are anchored to the inner surface of NE (see Fig. 1) *via* molecular interactions provided by a zoo of lamina-associated proteins such as LBR, LAP1, LAP2*β*, LEMD2, MAN1, PRR14, *etc*. (see, *e.g*., Refs. [7, 17, 30] and references therein). As such, given the length of LADs, it is intuitive to think that long strands of LADs bind to the inner-nuclear surface *via* multivalent molecular contacts, which can provide cumulative interaction energies on the order of ∼ 10 *k*_B_*T* [36, 37] easily, where *k*_B_ is the Boltzmann constants and *T* is the absolute temperature. This cumulative attraction strength can stabilize LADs strongly enough that the average distribution of peripheral chromatin may not change drastically during the lifetime of a healthy cell. Notably, a kinetic turnover between surface-attached and non-attached LAD segments can occur. This molecular picture can allow peripheral chromatin to be considered as an adsorbed polymer layer, which can reveal the functional role of peripheral chromatin structures.

In this study, we consider peripheral chromatin as a flexible polymer chain adsorbed onto the NE surface by employing de Gennes’ self-similar carpet model. Using scaling arguments, we estimate the thickness of the LAD layer and relate the polymer nature of that high-density chromatin layer to the experimentally observed steric hindrance of large molecules from the peripheral region of the nucleus. We relate our results and predictions to the available experimental data of nuclear chromatin distribution profiles, and lastly, discuss the implications and weaknesses of our model, together with future perspectives.

## II. SCALING THEORY OF PERIPHERAL CHROMATIN

During the interphase of eukaryotic cells, nuclear chromatin does not undergo microscopic changes. At this stage, individual chromosomes occupy distinct territories, but not all neighbor the nuclear boundary [2, 38]. This study only considers peripheral chromosomes, forming a straightforward polymer interface with the NE.

### A. Assumptions and justifications

In polymer physics, a Kuhn segment of size *b* = 2*𝓁*_p_ is defined as the largest segment of a polymer chain below which the polymer behaves as a rigid rod. The chemical composition of a polymer affects the persistence length 𝓁_p_, and thus, the Kuhn size *b*. However, polymer chains with distinctly different chemistry can exhibit universal conformational characteristics when considering chains composed of a sufficiently large number of Kuhn segments (*i.e*., *N* »1).

Considering the persistence length of chromatin on the order of 𝓁_p_ ∼ 1 kb to 10 kb [39–45], even the shortest reported LADs can correspond to *N*_L_ ∼ 10^1^ to ∼ 10^2^ Kuhn segments respectively. These values are much shorter than a single chromosome (*i.e*., with *N* ∼ 10^6^ Kuhn segments) but long enough to be statistically considered as a flexible polymer chain adsorbed on a surface.

Chromosomes are organized into chromatin fibers, which are protein-DNA complexes. One major protein complex organizing DNA into chromatin fiber is the histone octamer. Because the one-dimensional packing density of histone proteins varies along the fiber, chromatin is a highly heterogeneous polymer. Due to the variations in their histone packing densities [46, 47], heterochromatin and euchromatin can be structurally distinct and possibly have nonidentical persistence lengths [48]. For the same reason, values reported in the literature for chromatin persistence length vary largely, ranging between 𝓁_p_ ≈ 30 nm and 𝓁_p_ ≈ 300 nm [49–51]. Therefore, it is not straightforward to assign a clear difference between the persistence lengths of heterochromatin and euchromatin. Thus, we assume an identical Kuhn monomer size on the order of *b ≃* 10 nm as a lower limit for heterochromatin-rich LADs and euchromatin-rich inter-LADs. Note that throughout the article, ≃ signs denote scaling relations with neglected prefactors of order unity. Since scaling theories focus on the order magnitudes of the effects but not exact numbers, ignoring all prefactors on the order of unity, assuming equal Kuhn lengths for LADs and inter-LADs should not affect our scaling analysis. Hereon, by the term “*monomer*”, we refer to a “*Kuhn monomer*” of size *b*.

We assume that the chromatin in its native nuclear environment can be modeled as a polymer solution in the concentrated regime. That is, the volume fraction of chromatin is *ϕ ≳* 0.1. Single-particle tracking experiments reporting Rouse-type dynamics for the subsection of chromosomes support this assumption [52, 53].

We ignore the effects of electrostatic interactions since at physiological salt conditions (*i.e*., *c*_*s*_ ≈ 100 mM), chromatin could be treated as a neutral polymer for scaling purposes [49, 54]. In our calculations, there is also no selective attraction between chromatin segments that can lead to a phase separation.

Since the characteristic size of the chromosome is an order of magnitude smaller than the curvature radius of the nucleus (*i.e*., ∼ 10^0^ *μ*m *vs*. ∼ 10^1^ *μ*m), we treat the inner NE surface as planar.

To distinguish between a surface-adsorbed and a non-adsorbed chromatin monomer, the adsorption of LAD segments to NE is modeled under the “*weak adsorption*” approximation. The weak adsorption scheme also considers all weak interactions such as hydrogen bonding, ionic interactions, *etc*. Thus, instead of considering a subset of chromatin monomers anchored to NE by associated proteins with adsorption energies on the order of − *U* ≫ 1*k*_*B*_*T*, each chromatin monomer (whether LAD or inter-LAD) has an equal probability of interacting with the surface with an adsorption energy of *U* ≈ −*ϵ kT*, where 0 *< ϵ <* 1. LADs will be distinguished from inter-LADs *via* the formation of an adsorbed layer (see Sec. II C 1 below).

All parameters we consider are the ensemble averages of corresponding physical quantities. As argued above, an average number of Kuhn monomers per LAD and per inter-LAD (*N*_L_ and *N*_I_) are both considered to be much larger than unity. This regime of *long* polymers provides an excellent playground for scaling theories.

### B. Single-chain adsorption model for an isolated LAD

We begin our discussion by considering a single LAD composed of *N*_L_ monomers. While this discussion will provide insight for only an isolated LAD adsorbed on NE, it will help us introduce the concepts of chain adsorption, which will be used in the next section to describe the biologically relevant scenario of multiple independent LADs of a chromosome, overlappingly adsorbed on NE.

Consider an *n*-mer segment of a LAD, which is long enough to be considered as a flexible chain, but still much shorter than the whole LAD, *i.e*., 1 ≪ *n* ≪ *N*_L_. If the total attraction energy acting on all of its monomers is smaller than the thermal energy *k*_B_*T*, thermal fluctuations dominate over the cumulative surface attraction. As a result, the *n*-mer segment has an unperturbed configuration; its root-mean-square size, *r*(*n*) ≃ *bn*^*ν*^, is the same as that of an unperturbed/non-adsorbed *n*-mer. The exponent *ν* depends on the solvent quality, polymer volume fraction, and the molecular architecture of the polymer. In the biologically relevant concentrated regime of our concern, *ν* = 1*/*2 for linear polymers [55] and *ν <* 1*/*2 for cyclic polymers [56–67].

As *n* increases, the number of contacts that the *n*-mer segment can form with the surface, and hence, the effect of surface attraction on the segment, increases. At length scale

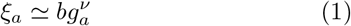

the cumulative adsorption energy of monomers in contact with the surface becomes on the order of thermal energy *k*_B_*T*. In Eq. (1), *ξ*_*a*_ is the size of an “*adsorption blob*” and *g*_*a*_ is the number of monomers per adsorption blob. The monomer volume fraction within an adsorption blob can be written as 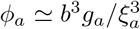. Using Eq. (1), we can write the volume fraction as a function of adsorption blob size as

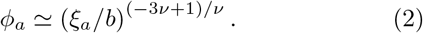

Thus, the number of monomers in contact with the surface inside an adsorption blob is 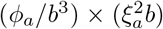. Here, *ϕ*_*a*_*/b*^3^ is the number density of monomers inside an adsorption blob, and 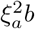 is the contact volume of the blob within a distance *b* to the surface. Thus, the adsorption energy of a blob is 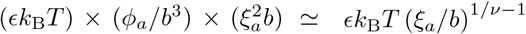. Equating this energy to thermal energy *k*_B_*T*, we obtain the size of an adsorption blob as [55]

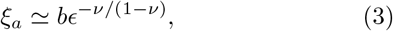

or the number of monomers in an adsorption blob as *g*_*a*_ ≃ *ϵ*^*−*1*/*(1*−ν*)^.

There are *N*_L_*/g*_*a*_ adsorption blobs along a single LAD. An isolated single LAD chain adsorbed on the nuclear periphery, therefore, behaves as a two-dimensional self-avoiding random walk of adsorption blobs on the surface (Fig. 2). Such a random walk of *N*_L_*/g*_*a*_ steps, each of size *ξ*_*a*_, has a root-mean-square end-to-end size of *R*_L_ ≃ *ξ*_*a*_ (*N*_L_*/g*_*a*_)^3*/*4^ [68], from which, we obtain

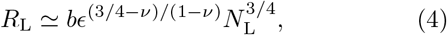

*via* Eq. (3). As the surface attraction becomes stronger with increasing *ϵ*, the adsorption blob size *ξ*_*a*_ shrinks, see Eq. (3), but the number *N*_L_*/g*_*a*_ of adsorption blobs per chain increases. The combined effect increases the two-dimensional end-to-end LAD size *R*_L_ (Fig. 2).

**FIG. 2.**
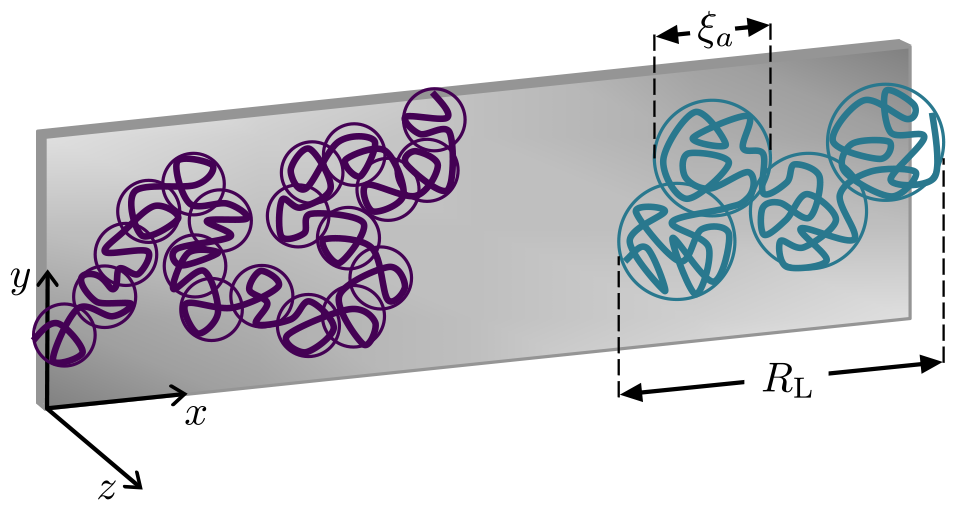
Illustration of two isolated chains adsorbed on a flat surface. Both polymer chains have the same number *N*L of Kuhn monomers of size *b*. The polymer on the left is adsorbed more strongly (with a larger ϵ) compared to the polymer on the right. For the more strongly adsorbed polymer on the left, number of monomers in an adsorption blob *g*_*a*_ and adsorption blob size *ξ*_*a*_ are smaller [see Eqs. (1) and (3)], while number of blobs *N*_L_*/g*_*a*_ and end-to-end size *R*_L_ are larger [see Eq. (4)].

Assuming a median value of *N*_L_ ≃ 10^2^ for a single LAD, we obtain *R*_L_ ≃ 100 nm for the size of a single LAD after assuming *ϵ ≃* 1. Considering a weaker attraction (*i.e*., *ϵ <* 1) decreases the LAD’s size since the segment loses contact with the surface (Fig. 2). Nevertheless, when the distance between neighboring adsorbed LADs is smaller than *R*_L_, those LADs interact by sterically repelling each other, and the above picture for isolated LADs loses validity. As will be discussed next, such a dense LAD environment alters polymer conformations and can lead to a steric environment for molecules diffusing through LADs.

### C. Multi-chain adsorption model for a single chromosome

Consider an individual chromosome in contact with NE located at *z* = 0, where *z* defines the distance from the surface toward the nuclear interior. A certain fraction of the chromosome monomers should be in contact with the NE surface to form LADs, while the rest can form extended loops (Fig. 1). Experiments show that only ∼ 30% of LADs identified by sequencing techniques are located at the nuclear periphery [7, 24, 69]. Thus, we can argue that there is insufficient space at the periphery to adsorb all LADs.

If LADs and inter-LADs are sufficiently long, we may treat them as independent chains. Accordingly, we suggest a molecular picture in which, near the nuclear periphery, many LADs compete for adsorption to the same limited surface space and sterically repel each other.

The adsorption energy is *ϵ k*_B_*T* per monomer in contact with NE. The cumulative surface energy of segments near the surface overcomes the steric repulsion between these segments, and consequently, monomer concentration near the surface is at its maximum. Contrarily, monomers positioned away from the surface have zero adsorption energy. As a result, the steric repulsion between segments becomes more dominant, favoring a less dense chromatin environment (Fig. 3a,c). Such a multi-chain adsorption scheme can be described by de Gennes’ self-similar carpet model [70], in which the competition between steric chain-chain repulsion and chain-surface attraction determines the properties of the structure of an adsorbed polymer layer.

**FIG. 3.**
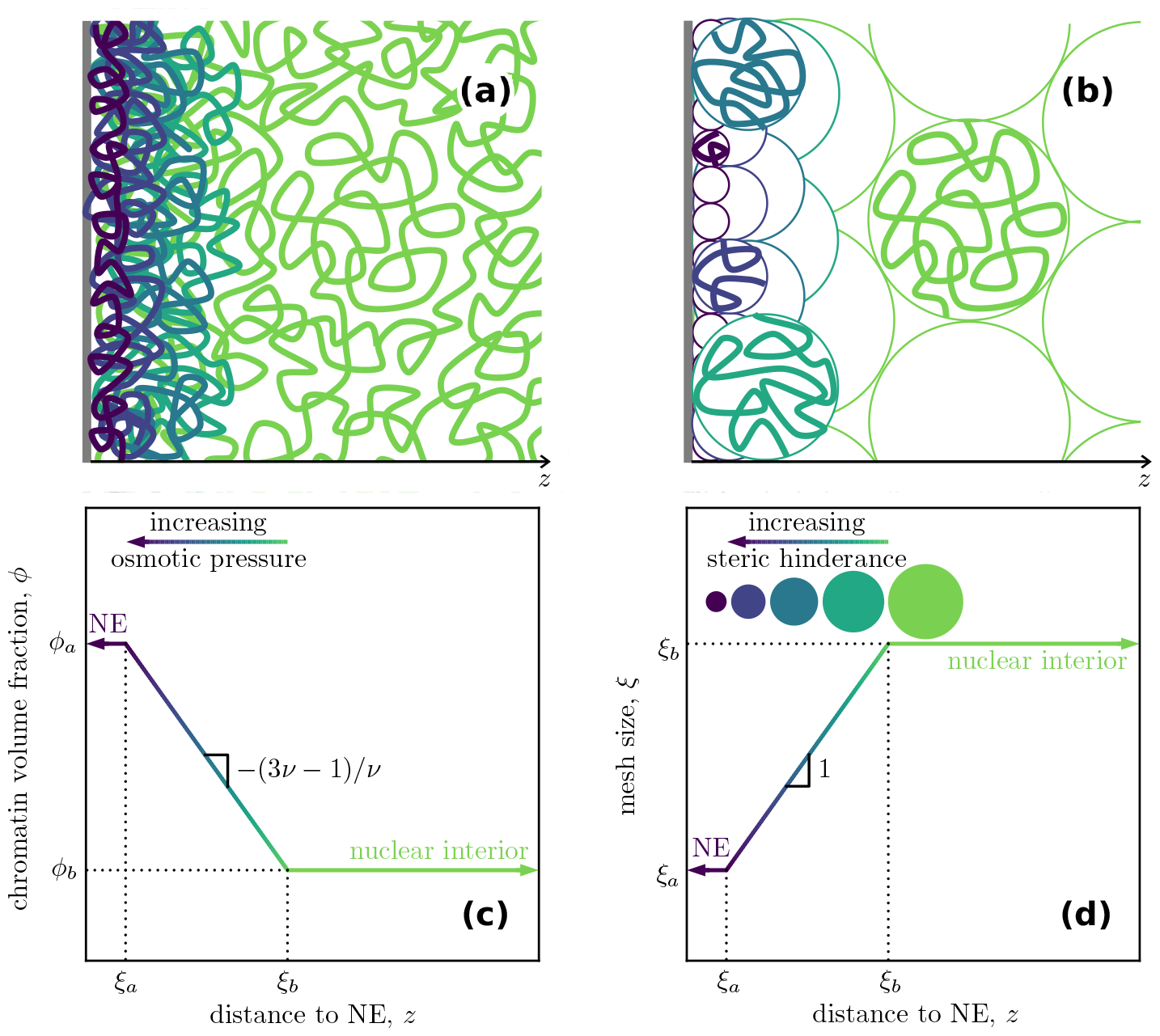
**(a)** and **(b)** Illustration of multi-chain adsorption on a flat surface. Osmotic pressure on chain segments is color-coded from high pressure close to the surface (purple) to low pressure away from the surface (green). In (b), only representative blobs are filled with chromatin segments for graphical simplicity. **(c)** and **(d)** The log-log plots of chromatin volume fraction *ϕ* and mesh (*i.e*., correlation blob) size *ξ* as functions of distance *z* from the adsorbing surface. As the chromatin density decreases and the mesh size approaches its bulk value, the steric hindrance of proteins decreases, as depicted by spheres of varying sizes in (d).

As discussed in the previous subsection, conformations of the adsorbed segments of this multi-chain structure are determined by the adsorption blobs of size *ξ*_*a*_ [see Eq. (3)]. The steric repulsion between chains defines a second length scale, the “*correlation blob*” size *ξ*, which is the average distance between segments of different chains (*i.e*., an effective mesh size). At the length scale of the correlation blob, the inter-chain steric repulsion energy is defined to be on the order of thermal energy *k*_B_*T*. Hence, at length scales below *ξ*, inter-chain steric repulsion is compensated by thermal energy, and chain segments have conformations unperturbed by the steric repulsion. That is to say, at length scales shorter than *ξ*, the conformation of a chain segment resembles that of an isolated chain. Thus, the correlation blob size can be written as *ξ ≃ bg*^*ν*^, where *g* is the number of monomers per blob.

The above definition of correlation blob size, *ξ*, gives the osmotic pressure (steric repulsion energy density) as *π ≃ k*_B_*T/ξ*^3^. At the surface, since the steric interactions (*k*_B_*T* per correlation blob) are compensated dominantly by the adsorption energy (*k*_B_*T* per adsorption blob), the correlation blob coincides with the adsorption blob (*i.e*., *ξ* = *ξ*_*a*_). The high osmotic pressure, due to the high density of monomers close to the surface, is relaxed gradually as the distance *z* to the NE surface increases. Such a relaxation increases correlation length *ξ* and decreases volume fraction *ϕ*. Inside the “*bulk*” of the nucleus, far away from NE, the correlation blob size *ξ*_*b*_ and the volume fraction *ϕ*_*b*_ are both independent of the distance *z*, and they are related by [55]

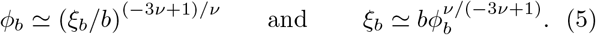

Eq. 5 also holds at an arbitrary distance *z* to the surface. The volume fraction of monomers inside a correlation blob is the same as the overall local volume fraction *ϕ*(*z*) at *z, i.e*.,

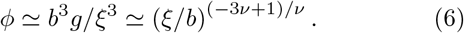

Hence, de Gennes’ self-similar carpet can be seen as layers of correlation blobs of increasing size *ξ*(*z*) with increasing distance *z* from the surface (Figs. 3b,d). The original model [70] argues a *z*-dependence for the correlation blob size, namely,

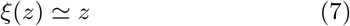

suggesting a linear relation between the correlation length and the distance *z* until it reaches the bulk value at *z* = *ξ*_*b*_. Plugging the above expression into Eq. (6) also provides a distance-dependent density profile:

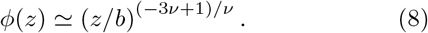

In summary, de Gennes’s self-similar carpet model predicts the chromatin volume fraction and the correlation blob size profiles between the nuclear boundary and interior as follows

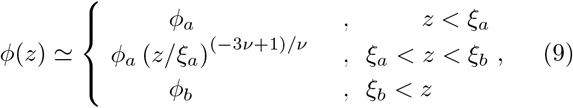

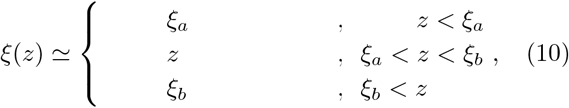

Eqs. 9 and 10 are plotted in Figs. 3c and d in log-log scales. Notably, the correlation blob size *ξ* is proportional to mesh size (*i.e*., the size sterically available to nuclear proteins). Thus, the smaller the *ξ* is, the stronger the steric hindrance near the nuclear boundary becomes. Fig. 3 highlights three distinct compartments of our model, each with a different chromatin density and mesh (correlation) size profiles. These three regions can be interpreted as follows:

#### 1. In close proximity of the nuclear envelope

At *z < ξ*_*a*_, the surface is covered with a layer of adsorption blobs of size *ξ*_*a*_. This high-density layer, which can be interpreted as a *true* LAD layer, has a thickness of *ξ*_*a*_, which is determined by the average size of adsorbed segments (Eq. (1)). Since some LAD segments cannot localize on the surface due to the limited surface area, they would be pushed towards the nuclear interior by the high chromatin-osmotic-pressure near the surface.

#### 2. In between the nuclear periphery and the nuclear bulk

As *z* increases along the interface region of *ξ*_*a*_ *< z < ξ*_*b*_, the high pressure and density at the nuclear periphery relaxes, and the chromatin density approaches its bulk value (Fig. 3c). The relaxation of surface effects increases correlation length *ξ* with increasing *z* (Fig. 3d). This increasing correlation size can allow large molecules to diffuse through the dense chromatin structure. In principle, layers of blobs in this interface region contain subsections of both LADs and inter-LADs.

#### 3. Inside the nuclear bulk

Far away from the nuclear periphery at *z > ξ*_*b*_, the abovementioned pressure relaxation is complete. The chromatin sections in this region do not *feel* the adsorbing surface from this distance. Here, the only effect of NE on the chromatin is the nuclear confinement. Experiments identified this interior region with low chromatin density and high transcriptional activity [4, 6, 25, 29]. This region is manifested by a surface-independent mesh size (correlation length) *ξ*_*b*_, which is larger than the mesh size near the surface.

## III. COMPARISON TO EXPERIMENTAL DATA FROM LITERATURE

To obtain a quantitative validation of our model, we facilitate the experimentally available data published in the literature. First, we compare our scaling predictions with the chromatin density profiles for eukaryotic cell types (Fig. 4 and Table I).

**TABLE I.**
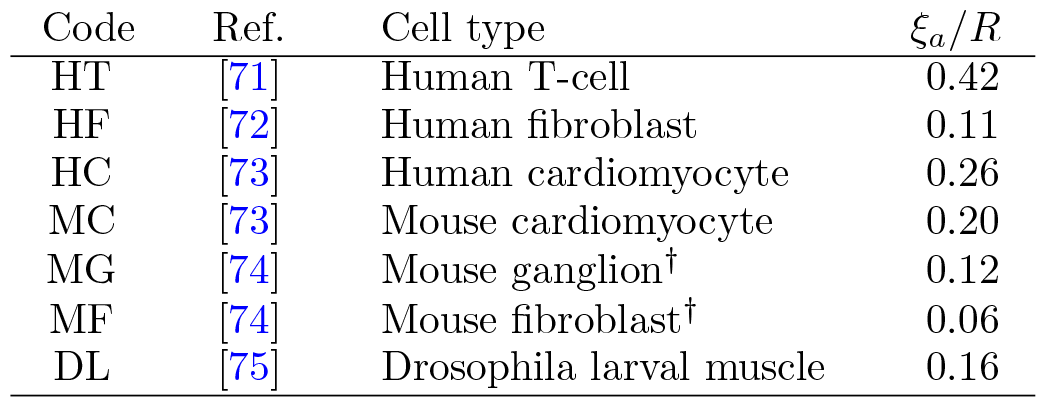
Fit parameters, *ξ*_*a*_*/R*, used to collapse the experimental data on a master curve in Fig. 4b. (^†^ Includes only gene-poor heterochromatin, but not the gene-rich euchromatin.)

| Code | Ref. | Cell type | $\xi_a/R$ |
| --- | --- | --- | --- |
| HT | [71] | Human T-cell | 0.42 |
| HF | [72] | Human fibroblast | 0.11 |
| HC | [73] | Human cardiomyocyte | 0.26 |
| MC | [73] | Mouse cardiomyocyte | 0.20 |
| MG | [74] | Mouse ganglion<sup>†</sup> | 0.12 |
| MF | [74] | Mouse fibroblast<sup>†</sup> | 0.06 |
| DL | [75] | Drosophila larval muscle | 0.16 |

**FIG. 4.**
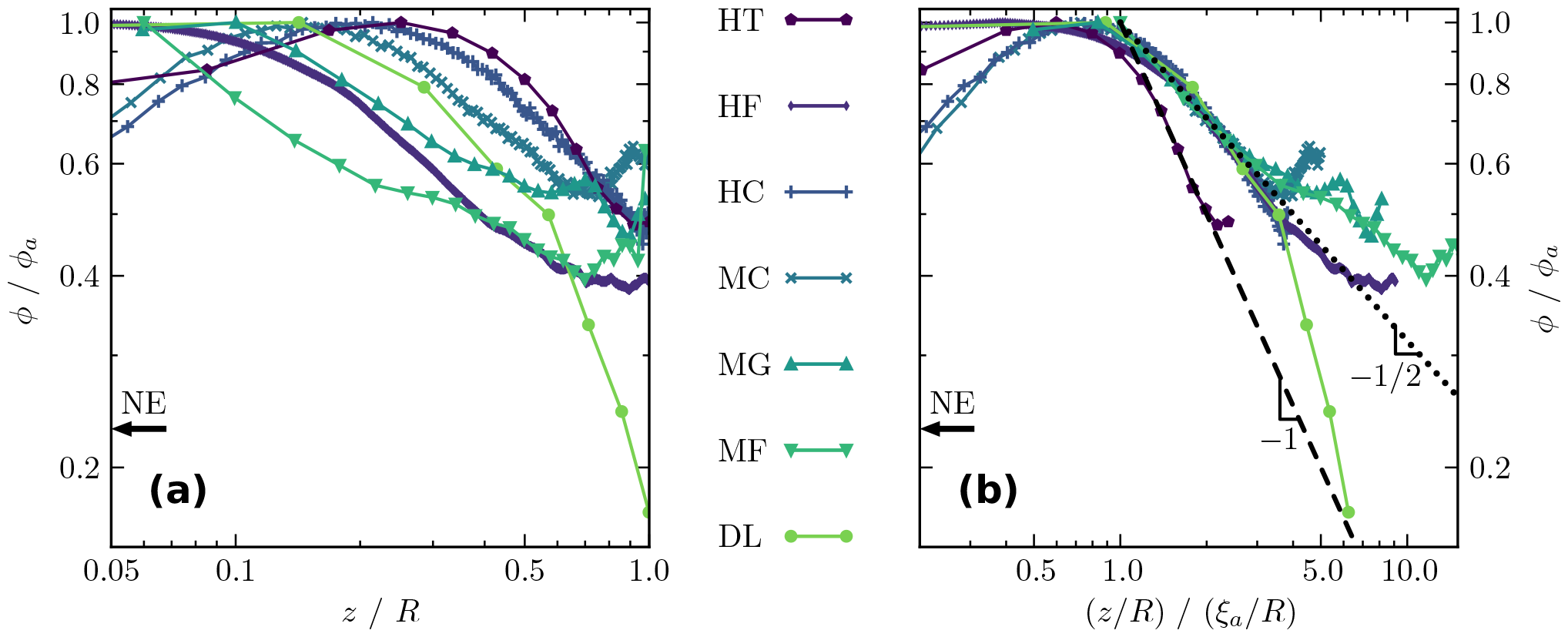
Log-log plots for the dependence of chromatin volume fraction *ϕ* on distance to nuclear periphery *z*. Data are from various eukaryotic cell types; see Table I for abbreviations. In both panels, *ϕ* data is scaled by the maximum measured volume fraction *ϕ*_*a*_ for that cell type. In the left panel, *z* is scaled by cell radius *R*, such that *z/R* = 1 is at the nuclear center. The decrease in chromatin density close to the nuclear periphery at small *z/R*, is due to a depletion region not considered in the theory. In the right panel, *z/R* is scaled with *ξ*_*a*_*/R* to obtain a data collapse in accordance with Eq. (11). Fitting parameteres for *ξ*_*a*_*/R* are given in Table I. The dashed and dotted lines show our predictions of *ϕ/ϕ*_*a ≃*_ (*z/ξ*_*a*_)^−1^ and *ϕ/ϕ*_*a ≃*_ (*z/ξ*_*a*_)^−1*/*2^; see Fig. 3c and Eq. (11) with *ν* = 1*/*2 and *ν* = 2*/*5 respectively. The data points are joined to guide the eye.

In literature, the nuclear chromatin density profiles were often reported in the units of fluorescence intensity or arbitrary units as a function of nuclear radial distance (*i.e*., radial position scaled by the nuclear radius *R*). Therefore, to compare our results summarized in Eq. (9) with such data, we normalize the density axis by the maximum value of the signal for the corresponding cell type. The maximum value is matched to the density value near the surface in our model (*i.e*., *ϕ*_*a*_). Then, we compare the rescaled data with our normalized density distribution (*i.e*., *ϕ/ϕ*_*a*_) (Fig. 4b).

In general, as shown in Fig. 4a, decreasing chromatin concentration with increasing distance from the nuclear periphery is a common hallmark of the density profiles in all simulations and experiments [35, 71–79] and agrees well with our model, see Eq. (9) and Fig. 3c. This can be seen quantitatively in the master curve combining all the available data by using the expression

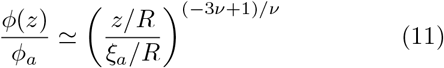

derived from Eq. (9) for *ξ*_*a*_ *< z < ξ*_*b*_. To obtain the overlap in Fig. 4b, we use *ξ*_*a*_*/R* as a free fit parameter in Eq. (11) for all cell types (see Table I for the fit values). We observe that the thickness of the LAD layer relative to the nuclear radius is in the range of about *ξ*_*a*_*/R ≃* 0.1 − 0.2, which agrees well with high-resolution microscopy images [24, 31, 69]. The only exception is the human T-cell, for which we predict a *ξ*_*a*_*/R* value of about 40% (Table I). Also, note that data for this cell type is singled out from the other collapsed data sets in Fig. 4b.

The data in Fig. 4b highlights a concentration decay from the nuclear surface to the bulk. However, for the human T-cell, this decay is stronger; while our model predicts *ϕ/ϕ*_*a*_ = (*z/ξ*_*a*_)^*−*1^ for the human T-cell with an exponent of *ν* = 1*/*2, all other data can be collapsed onto a universal curve of *ϕ/ϕ*_*a*_ = (*z/ξ*_*a*_)^*−*1*/*2^ away from the nuclear center and periphery. This expression is obtained if an exponent of *ν* = 2*/*5 is chosen for the statistical conformations of the LAD and inter-LAD segments; see Eq. (11). The exponent *ν* = 2*/*5 describes the data relatively well and is consistent with the exponent suggested for cyclic polymers [56–60]. However, it is larger than other predictions for cyclic chains, *e.g*., *ν* = 1*/*3 [61, 62]. This discrepancy could reflect a crossover window from a *Gaussian regime* of *ν* = 1*/*2 for short cyclic chains to a *compact regime* of *ν* = 1*/*3 for longer cyclic polymers. Indeed, an exponent of *ν* = 2*/*5 is observed in simulations for a wide crossover regime [63–67]. In that sense, the outlier data set in Fig. 4b, human T-cells, can be rationalized by the argument that the peripheral chromatin in human T-cells is in the Gaussian regime.

Next, we compare the correlation lengths that we predict with the available experimental data reporting the effective porosity of nuclear chromatin. By microinjecting fluorescent-labeled dextran probe molecules of varying molecular mass *M* and size *d* into interphase HeLa nuclei, Görisch *et al*. correlated the nuclear distribution of the particles with the nuclear porosity near and away from the periphery [35]. While small dextran molecules (*i.e*., *d ⪅* 20 nm) with *M* ≤ 77 kDa can diffuse close to the nuclear periphery, larger molecules cannot. For those larger molecules, the dextran density increases away from the NE and levels off inside the nuclear bulk.

This finding is in qualitative agreement with our predictions for the mesh size [see Eq. (6)]

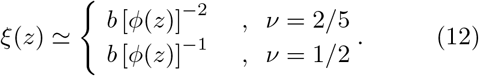

Note that the above predictions are much stronger than the inverse square root dependence obtained by considering chromatin as a hexagonal packed crystal [80].

The smallest length scale in our model is the Kuhn length (or the persistence diameter), *b* = 2𝓁_*p*_. Such a scaling model can only make limited predictions about the phenomena related to length scales smaller than *b*. One such prediction is that, since the Kuhn length defines the size of the smallest loops that can be formed by the polymer chain inside the dense chromatin environment, any molecule smaller than *b* can find pores to diffuse along. This simple picture is adequate to describe the smallest mesh size of *b* ≈ 20 nm observed by Görisch *et al*. close to the nuclear periphery. In fact, this is on the order of the Kuhn size of chromatin (*i.e*., *b* ≃ 10 nm).

A second steric barrier observed in the experiments is the mesh size of *ξ* ≈ 60 nm, which can be inferred as the size *ξ*_*a*_ of the adsorption blobs of our model [35, 80]. For the HeLa nuclei of radius *R* ≈ 1 *μ*m, this adsorption blob size value leads to *ξ*_*a*_*/R* ≈ 0.06, in accord with the range of values presented in Table I. We note that by plugging the values *ξ*_*a*_ ≈ 60 nm and *b* ≈ 20 nm into Eq. (1), we obtain the number of Kuhn monomers in an adsorption blob as *g*_*a*_ *≃* 10 for both *ν* = 1*/*2 and *ν* = 2*/*5 exponents. Recalling that our scaling theory assumes *g*_*a*_ ≫ 1, a value of *g*_*a*_ *≃* 10 is at the limiting boundary of the validity of our model.

Finally, from the relative thickness *ξ*_*a*_*/R*, we predict the relative nuclear volume of the LAD layer as

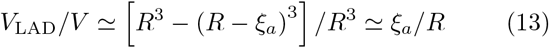

where *V* ≃ *R*^3^ is the nuclear volume. For a nuclear radius of around *R* ≈ 5 *μ*m, using the values in Table I, we obtain a relative volume of the LAD layer in the range *V/V*_LAD_ ≃ 10 − 20%, with an average LAD layer thickness of *ξ*_*a*_ *<* 1 *μ*m. This prediction agrees well with high-resolution microscopy images [24, 31, 69].

## IV. DISSCUSSION

By revisiting de Gennes’ self-similar carpet model, we argue that lamina-associated chromatin domains can be considered as multiple independent chain segments (of a chromosome) attracted to a surface with an energy of *U* = − *ϵ k*_B_*T* per adsorbed Kuhn monomer. In our calculations, we assume the chromatin layer adjacent to the surface with a thickness *ξ*_*a*_ is all occupied by LADs.

The polymer model considered here does not separate LADs and inter-LADs with respect to their chemical or structural properties. Yet, one can conclude that chromatin sections away from the nuclear periphery are of inter-LAD nature implicitly. Our multi-chain model assumes that the layer closest to the nuclear periphery is filled almost entirely with LADs, and implicitly, chromatin sections away from the nuclear periphery are of inter-LAD nature. In principle, the LAD layer can also contain a substantial amount of inter-LADs, especially if the fraction of LADs is small (*i.e*., *N*_L_ ≪ *N*_I_). However, DamID and ChIP-seq experiments associate ∼ 40% of the genome as LADs [7, 9, 81], suggesting a dominant LAD presence within the adsorption layer. Note that, in a synthetic system of a copolymer with *sticky* and *non-sticky* domains, decreasing the fraction of sticky domains would affect the polymer structure near the confining surface.

The scaling approach we present here does not distinguish between chromatin sections permanently tethered to NE and those interacting weakly with the periphery. Nevertheless, the statistical nature of the scaling analyses favors surface localization limited by steric repulsion between LAD segments. Thus, the detailed nature of the surface localization does not change the outcome and provides relatively good chromatin distributions as compared to experiments. However, distinguishing tethering from weak attraction or varying the chain stiffness affected the contact probabilities at ∼ 100 kb scale [45], but averaging over many segments in contact with NE can provide a picture similar to our predictions. It is also intuitive to think that more permanent contacts with NE can give LADs structural properties such as those observed in cyclic polymer brushes [82–84]. In fact, assuming a cyclic architecture for an inter-LAD bordered by two LADs, protruding towards the nuclear interior, forming loops, describes the chromatin density profile data for various eukaryotic cell types relatively well (Fig. 4). In that sense, the stronger decay of chromatin volume fraction for the outlier data set in Fig. 4b (human T-cells) could be due to the T-cell chromatin being in the Gaussian polymer regime. It might also be due to the constrained release of lamina-associated enhancers and genes from the nuclear envelope, as observed during Jurkat T-cell activation [85], which changes the 3d organization of the genome.

Our model explains the steric hindrance of large molecules from the nuclear periphery. Adsorption of LADs to NE generates a distinct structure of chromatin with a mesh size smaller than that in the bulk of the nucleus (*i.e*., *ξ < ξ*_*b*_). This picture leads to an effective hindrance mechanism. The smallest mesh size is *ξ*_*a*_ at the proximity of NE. The same *ξ*_*a*_ defines the thickness of the densest layer on the surface, *i.e*., the thickness of the LAD layer. As the distance to NE increases, the mesh size grows towards that inside the nuclear bulk (*ξ*_*b*_). This hindrance can exclude large proteins of transcription machinery from the LADs and reinforce the suppressive environment of LADs *via* this non-specific effect that does not depend on the nucleic acid sequence. The exact mechanism can allow larger mesh sizes, for instance, near the nuclear pore complexes around which euchromatin is relatively sparse, allowing more transcriptional activity near LADs. The hindrance of very small molecules (smaller than the persistence diameter of chromatin, ∼ 20 nm) cannot be explained by the mechanism of growing mesh size *ξ* but could be due to the coalescence of nucleosomes into an effective melt similar to polymer brushes [86, 87]. Consistently, a recent study [88] discusses that surface-attached LADs have less gene activity than non-attached LADs, which reside at the interface between the so-called “compartment A” and “compartment B”. This result supports the increasing mesh size along the interface region. Proteins regulating the genetic machinery cannot reach the dense layer of surface-attached LADs, while they reach the non-attached LAD segments of the larger mesh size.

The polymer model considered here does not separate heterochromatin and euchromatin. A more detailed model can separate these two chromatin types, possibly considering copolymers with two different persistence lengths [48] or surface-affinities. Nevertheless, we anticipate that within the limits of valid biological parameters, decreasing steric hindrance and decreasing chromatin density towards the nuclear interior will be preserved even in a more complex polymer model. This is also evident from the experimental data in which more compact chromocenter regions of the genome have a mild effect on chromatin distribution, albeit these faces cause higher heterochromatin concentration near their surface. (See, *e.g*., MG and MF data in Fig. 4b and the corresponding Ref. [74].)

An interesting direction to explore would be to investigate the structural support provided by the dense LAD layer to NE [33]. It is experimentally demonstrated that the loss of the adsorption layer (due to the repression of proteins that bind chromatin to NE) results in deformed nuclear morphology in various diseases [15, 17, 19].

Overall, our theoretical approach demonstrates that the polymer nature of the genome can generate functional properties such as non-specific steric hindrance of transcriptional components in the nucleus. Various polymer models, which are well studied in the context of synthetic polymers, can reveal the structural complexity of genome organization, and hence, can help us extend our knowledge of the relationship between 3d chromosome organization, genetic regulation, and nuclear morphology.

## ACKNOWLEDGMENTS

We thank Jaroslaw Paturej for the valuable discussions. This work has been supported by the National Science Center, Poland (Grant Polonez Bis No. 2021/43/P/ST3/01833) and The Council of Science and Technology of Turkey (TUBITAK) Grant no 122F309.

